# Loosenin-like proteins from *Phanerochaete carnosa* impact both cellulose and chitin fiber networks

**DOI:** 10.1101/2022.07.01.498415

**Authors:** Mareike Monschein, Eleni Ioannou, Leamon AKM AL Amin, Jutta J. Varis, Edward R. Wagner, Kirsi S. Mikkonen, Daniel J. Cosgrove, Emma R. Master

**Affiliations:** Department of Bioproducts and Biosystems, Aalto University, Kemistintie 1, 02150 Espoo, Finland; Department of Food and Nutrition, University of Helsinki, Agnes Sjöbergin katu 2, 00014, Helsinki, Finland; Department of Biology and Center for Lignocellulose Structure and Formation, 208 Mueller Laboratory, Pennsylvania State University, University Park, State College, PA 16802, United States; Department of Chemical Engineering and Applied Chemistry, University of Toronto, 200 College Street, Toronto, Ontario, M5S 3E5, Canada

**Keywords:** loosenin, expansin, *Phanerochaete carnosa*, cellulose, chitin

## Abstract

Microbial expansin-related proteins are ubiquitous across bacterial and fungal organisms, and reportedly play a role in the modification and deconstruction of cell wall polysaccharides including lignocellulose. So far, very few microbial expansin related proteins, including loosenins and loosenin-like (LOOL) proteins, have been functionally characterized. Herein, four LOOLs encoded by *Phanerochaete carnosa* and belonging to different subfamilies (i.e., PcaLOOL7 and PcaLOOL9 from subfamily A; PcaLOOL2 and PcaLOOL12 from subfamily B) were recombinantly produced and the purified proteins were characterized using diverse cellulose and chitin substrates. Whereas all of the purified PcaLOOLs weakened cellulose filter paper and cellulose nanofibril networks (CNF), none significantly boosted cellulase activity on the selected cellulose substrates (Avicel and Whatman paper). Binding of PcaLOOLs to alpha-chitin was higher than to cellulose (Avicel), and highest at pH 5.0. Notably, whereas PcaLOOL9 reduced the yield strain of chitin nanofibrils (ChNF) in a protein-dose dependent manner, the reverse pattern was observed for PcaLOOL7 despite belonging to the same LOOL subfamily. The current study reveals the potential of microbial expansin-related proteins to impact both cellulose and chitin networks, and provides further evidence pointing to a non-lytic mode of action.

## 1. Introduction

Expansins were first identified through studies of acid-induced growth of cucumber hypocotyls (*Cucumis sativus*), where they trigger cell wall loosening without evidence of lytic activity (1-3). Plant expansins have since been implicated in fruit ripening, root hair elongation, germination, and pollen tube penetration and other developmental processes (4,5). Although their mode of action remains elusive, detailed biochemical and biophysical studies indicate that expansins disrupt noncovalent bonds at so-called ‘biomechanical hotspots’, which are load-bearing junctions between cellulose microfibrils or between cellulose microfibrils and other matrix polymers (6-8). Expansins have been identified across land plants (4, 5, 9, 10) and the ubiquity of expansin-related proteins among microorganisms has since been verified through genome sequencing (9, 11-14).

All expansins exhibit a distinctive two-domain structure of 250 to 275 amino acids. The N-terminal domain (D1) is a six-stranded double-psi beta-barrel (DPBB) which is packed tightly next to a C-terminal domain (D2) having a β-sandwich fold (15). D1 and D2 domains align to form a long, shallow groove with highly conserved polar and aromatic residues suitable to bind a twisted polysaccharide chain (15-17). D1 is structurally related to the catalytic domain of family-45 glycoside hydrolases (GH45); however, it lacks the aspartate that serves as the catalytic base in GH45 enzymes (16) and so far, no lytic activity has been observed for any plant expansin or bacterial expansin-like protein (10). D2 is homologous to group-2 grass pollen allergens and has been classified as a family-63 carbohydrate binding module (CBM63) (17).

Expansins and expansin-related proteins are classified based on phylogenetic analysis (13, 14, 18). Plant-derived expansins are classified as α-expansins (EXPA), β-expansins (EXPB), and two small groups of expansin-like sequences (EXLA and EXLB). Microbial expansins (EXLX) include proteins with both D1 and D2 domains. Some expansin-related proteins from microbes possess other domain architectures, including proteins comprising only the D1 domain (e.g., loosenins (19), cerato-platanins (20)) and multidomain proteins comprising other domains in addition to D1 and D2 (e.g., swollenins (16, 21, 22). So far, most studies of microbial expansin-related proteins have focused on their potential to boost enzymatic hydrolysis of lignocellulosic substrates to sugars (23, 24, 25, 26). Reported impacts on lignocellulose deconstruction, however, vary and depend strongly on the biomass source (27).

In an effort to deepen our understanding of the molecular function and applied potential of microbial expansin-related proteins, we recombinantly produced and characterized four loosenin-like (LOOL) proteins encoded by *Phanerochaete carnosa* and expressed during growth on wood substrates (28, 29). Only two loosenins were previously characterized: LOOS1 from the white-rot basidiomycete *Bjerkandera adusta* (19) and N2 of *Neurospora crassa* (30). Both LOOS1 and N2 reportedly disrupt the structure of cotton fibers (19, 30); pretreatment of *Agave tequilana* fibers with LOOS1 also increased the susceptibility of the material towards enzymatic hydrolysis, enhancing the rate of reducing sugar release by up to 7.5-times (19). The PcaLOOLs characterized herein weakened cellulose filter paper and altered the rheological properties of cellulose and chitin nanofibers; however, these impacts did not translate to a substantial change in the enzymatic saccharication of isolated cellulose and chitin substrates.

## 2. Materials and methods

### 2.1 Substrates and chemicals

Cellulose nanofibers (CNF) were prepared from never-dried bleached Kraft birch pulp by fluidizing in deionized water six times (six passes) using a Voith LR40 homogenizer, as desribed in Österberg et al. (2013) (31). The resulting CNF samples with a solid content of 2.8% w/v were stored at 4°C. Chitin nanofibers (ChNF) were extracted from crab shells and then purified and mechanically disintegrated as described in Liu et al. (2019) (32). The resulting ChNF samples with a solid content of 1.1% w/v were stored at room temperature. Avicel® PH-101 (∼50 µm particle size; Cat.# 11365), xylan from birch wood (Cat.# 95588), Whatman® qualitative filter paper grade 1 (Cat.# WHA1001185), Whatman® qualitative filter paper grade 3 (Cat.# WHA1003185), and peptidoglycan from *Micrococcus luteus* (Cat.# 53243), were purchased from Sigma-Aldrich (St. Louis, MO, USA). Chitin from shrimp shells (Cat.# P-CHITN), glucomannan (Konjac; low viscosity; Cat.# P-GLCML), carboxymethyl cellulose 4M (CMC; Cat.# P-CMC4M), xyloglucan (Tamarind; Cat.# P-XYGLN), hexaacetyl-chitohexaose (Cat.# O-CHI6), cellohexaose (Cat.# O-CHE), and β-glucan (yeast; alkali soluble; Cat.# P-BGYST) were obtained from Megazyme (Bray, Ireland). Thermo Fisher Scientific (Waltham, MA, USA) was the supplier of Pierce™ BCA Protein Assay Kit (Cat.# 23225). All other chemicals were reagent grade.

### 2.2 Enzymes

Cellulase mixture Cellic®CTec2 (cellulase, enzyme blend; Cat.# SAE0020), Celluclast® (cellulase from *Trichoderma reesei*; Cat.# C2730), chitinase from *Trichoderma viride* (Cat.# C8241), lysozyme (from hen egg white; Cat.# 10837059001) and α-chymotrypsin (from bovine pancreas; Cat.# C6423) were obtained from Sigma Aldrich; PNGase F from New England Biolabs (Ipswich, MA, USA; Cat.# P0705S); *endo*-1,4-β-D-glucanase (cellulase from *Trichoderma longibrachiatum*, glycoside hydrolase family 7; Cat.# E-CELTR) from Megazyme. BsEXLX1 (expansin from *Bacillus subtilis*) was recombinantly expressed in *Escherichia coli* BL21 and purified as described in Georgelis et al. (2011) (17).

### 2.3 Sequence analysis

Amino acid sequences of loosenin-like proteins PcaLOOL2 (GenBank code EKM55357.1), PcaLOOL7 (GenBank code EKM53490.1), PcaLOOL9 (GeneBank code EKM52742.1) and PcaLOOL12 (GenBank code EKM51974.1) from *P. carnosa* (28) were retrieved from the NCBI protein database (http://www.ncbi.nlm.nih.gov/protein/). Secretion signal prediction was performed using the SignalP v. 4.1 web server (33). Amino acid sequences were aligned with Clustal Omega (34) and sequence similarities were visualized using the program ESPript 3 (http://espript.ibcp.fr) (35). Expasy tools ProtParam (36) and Prosite (37) were used to compute physico-chemical protein parameters and predict glycosylation sites; DISULFIND (38) for the detection of disulphide bridges.

### 2.4 PcaLOOL production and purification

Selected PcaLOOLs were expressed in *Pichia pastoris* strain SMD1168H following the manufacturer’s instructions (Invitrogen™, Thermo Fisher Scientific). Briefly, codon optimized genes encoding each PcaLOOL were obtained as subcloned in pPICZαA plasmids with a C-terminal 6 x His tag (GenScript, Piscataway, NJ, USA). *P. pastoris* transformants were screened for protein expression by immuno-colony blot using nitrocellulose membranes (Bio-Rad Laboratories, Hercules, CA, USA; Cat.# 1620115), Tetra·His antibodies (Qiagen, Hilden, Germany; Cat.# 34670), Anti-Mouse IgG (whole molecule), peroxidase antibodies produced in rabbit (Sigma-Aldrich; Cat.# A9044), and SuperSignal™ West Pico PLUS chemiluminescent substrate (Thermo Fisher Scientific; Cat.# 34579). Precultures (4 × 50 mL in 500 mL baffled flasks) of the best transformant for each PcaLOOL were then grown in buffered glycerol-complex medium (BMGY; 100 mM potassium phosphate buffer (pH 6.0), 2% (w/v) peptone, 1% (w/v) yeast extract, 1.34% (w/v) yeast nitrogen base, 4×10^−5^% (w/v) biotin, 1% (v/v) glycerol) at 30°C and 100 rpm until an OD_600_ of ∼ 6 was reached. Cells were harvested by centrifugation at 15°C, 1500 x g for 10 min and the obtained pellets were suspended in 4 × 300 mL methanol-complex medium (BMMY) containing 0.5% (v/v) methanol instead of glycerol. Each 300 mL cultivation was performed in a 2.5 L Tunair® shake flask covered with two layers of sterile Miracloth. Methanol was added to 1% (v/v) every 12 h and induction was continued over 132 h at 28°C and 130 rpm. After the induction, culture supernatants were recovered, filtered and the secreted recombinant proteins were purified by affinity chromatography using Ni-NTA resin (Qiagen; Cat.# 30230). Specifically, supernatants were concentrated using a Vivaflow® 200 crossflow cassette with a 10000 MWCO PES membrane (Sartorius, Göttingen, Germany; Cat.# VF20P0) and loaded onto a 5 mL GE Healthcare HisTrap™ FF Crude prepacked column (Thermo Fisher Scientific; Cat.# 11723219). Proteins were eluted by FPLC using an ÄKTA purifier (Amersham BioScience, Amersham, UK).

The purified proteins were concentrated and exchanged to 10 mM sodium citrate buffer (pH 5.0) using Vivaspin® Turbo 4 ultrafiltration units with a 5000 MWCO PES membrane (Sartorius; Cat.# VS04T11). The purity and concentration of each PcaLOOL was assessed by SDS-PAGE and the Pierce™ BCA Protein Assay Kit, respectively, prior to storage at −80°C. To confirm protein identity, deglycosylated forms of the purified PcaLOOLs were prepared using PNGase prior to digestion with α-chymotrypsin and analysis using an ultrafleXtreme MALDI -TOF/ TOF mass spectrometer (Bruker, Billerica, MA, USA).

### 2.5 Circular dichroism (CD) spectroscopy

The recombinant PcaLOOLs were diluted to 0.1 mg/ mL in H_2_O. CD spectroscopy was performed using a Chirascan CD spectrometer (Applied Photophysics, Leatherhead, UK). CD data were collected between 190 to 280 nm at 22°C using a 0.1 cm path-length quartz cuvette. CD measurements were acquired every 1 nm with 0.5 s as an integration time and repeated three times with baseline correction. Chirascan Pro-Data Viewer (Applied Photophysics) was used to convert direct CD measurements (θ; mdeg) to mean residue molar ellipticity ([θ]MR), and secondary structures were predicted using the BeStSel web server (39) from 190 to 250 nm and a scale factor of 1. Thermal unfolding was recorded from 20°C to 80°C between 190 and 280 nm with a 2°C step size at 1°C/ min ramp rate with ± 0.2°C tolerance. The melting temperature was analyzed with Global3 (Applied Photophysics).

### 2.6 Test for hydrolytic activity

Purified proteins were tested for hydrolytic activities towards xylan from birchwood, CMC and glucomannan. Substrates were suspended in 50 mM sodium acetate buffer (pH 5.0) to a final concentration of 1% (w/v); 125 µL of each substrate were then transferred to separate wells in a 96-well plate (Thermo Fisher Scientific) and supplemented with 0.01 mg/ mL PcaLOOLs and milliQ water to a final sample volume of 250 µL. BSA was used as a reference. Plates were incubated for 16 h in a ThermoMixer®C set at 40°C and 700 rpm. Reducing sugar concentration was determined by the para-hydroxybenzoic acid hydrazide (PAHBAH) assay calibrated against glucose (40).

### 2.7 Test for lytic activity

For the turbidimetric assay, water insoluble substrates (β-glucan from yeast, peptidoglycan from *Micrococcus luteus*) were suspended at 0.35 mg/ mL in 50 mM sodium acetate buffer (pH 5.0) and supplemented with 0.05 mg/ mL PcaLOOL or BSA. Reactions were incubated in a ThermoMixer® C set at 1000 rpm and 25°C for 0 – 24 h. At regular intervals, 0.12 mL samples were collected and analysed spectrophotometrically at 600 nm to detect substrate solubilization. Samples without protein served as negative controls.

For the analysis of reaction samples by thin layer chromatography (TLC) polysaccharide substrates (xyloglycan, peptidoglycan, β-glucan) were suspended at 0.35 mg/ mL in 50 mM sodium acetate buffer (pH 5.0) and supplemented with 0.05 mg/ mL PcaLOOL. Oligosaccharide substrates (cellohexaose, hexaacetyl-chitohexaose) were suspended at 1 mM in 50 mM sodium acetate buffer (pH 5.0) and mixed with 0.125 mg/ mL PcaLOOL. Reactions were incubated as described above for a period of 24 h. BSA was used as a reference on polysaccharide and oligosaccharide substrates; lysozyme and *endo*-1,4-β-D-glucanase were also included as reference treatments of oligosaccharide substrates. Samples (2 µL of polysaccharide reactions, 10 µL of oligosaccharide reactions) and standards (1 µL of 250 mM glucose, 250 mM xylose or 225 mM N-acetylglucosamine) were applied to a TLC Silica gel 60 F254 (Sigma-Aldrich).

Chromatograms were developed in n-propanol/ 25% ammonia (2:1) solvent mixture as eluent. Spots were visualized by spraying with 10% sulfuric acid in ethanol followed by heating of the plates using a Steinel HL 1920E hot air blower.

### 2.8 Binding studies

Protein adsorption to different substrates was monitored by a pull-down assay. Specifically, 2.5 mg/ mL of cellulosic substrate (Avicel® PH-101 or Whatman® qualitative filter paper grade 1) were weighed into 1.5 mL Eppendorf® LoBind microcentrifuge tubes (Sigma-Aldrich) and suspended in a final reaction volume of 500 µL 50 mM sodium acetate buffer (pH 5.0) supplemented with 0.1 mg/ mL target protein. Bovine serum albumin (BSA) (Sigma-Aldrich; Cat.# A3059) was used as a reference. Protein blanks were prepared using protein solutions without substrate; substrate blanks contained substrate suspensions without protein. Samples were prepared in triplicate and incubated for 1 h at room temperature on a tube rotator set to 20 rpm; supernatants were then recovered by centrifugation (15000 rpm for 10 min) and protein concentrations were measured using the Pierce™ BCA Protein Assay Kit.

PcaLOOL binding to chitin from shrimp shells was performed as described above, except at different pH values: 50 mM sodium citrate (pH 3.5); 50 mM sodium acetate buffer (pH 5.0), 50 mM sodium phosphate (pH 6.0), and 50 mM sodium phosphate (pH 7.0).

### 2.9 Paper weakening assay

The ability to weaken filter paper was analysed as described in Cosgrove et al. (2017) (41). Briefly, Whatman® qualitative filter paper grade 3 was cut into 10 × 2.0 mm strips and soaked in 1 mL of 20 mM MES buffer (pH 6.0) containing 0.2 mg/ mL protein. BsEXLX1 was used as a positive reference. Incubation was performed with gentle inversion to equilibrate the solution with the strips at 25°C for 4 h. After incubation, filter paper strips were fixed between two clamps of a custom-built extensometer (41) and extended at 1.5 mm/ min while recording the tensile force on a digital chart recorder. The maximum force attained was taken as the breaking force.

### 2.10 Cell wall extension (creep) assay

Etiolated wheat (*Triticum aestivum* L. cv Pennmore) coleoptiles were prepared as described in Cosgrove et al. (2017) (41), heat inactivated by a 15-s dip in boiling water and fixed between two clamps of a custom-built constant-force extensometer for cell wall creep experiments (1), which resulted in a tensile force of 20 g on the specimen. The specimen was kept in 200 µL 20 mM MES buffer (pH 6.0) for 10 min, before the buffer was exchanged for 50 mM NaOH to increase the sensitivity of the material in the creep assay. After 10 min, coleoptiles were rinsed in 20 mM MES buffer (pH 6.0) and after an additional 15 min the buffer was replaced with fresh buffer containing 0.2 mg/ mL protein.

### 2.11 Rheological measurements

Rheological measurements were performed using cellulose nanofibers (CNF) and chitin nanofibers (ChNF). For CNF, the stock solution (2.8% w/v) was first dispersed in 0.423 mL H_2_O and sonicated with a tip sonicator (Q500; QSonica, Newton, CT, USA) at 20% amplitude for 2 min (2 s on/ off cycles). For both CNF and ChNF treatments, final concentrations of sample components were 0.6% substrate, 5 mM sodium acetate buffer (pH 3.5), and between 0.25 - 2 mg/ mL of protein. Treatments with CNF were also performed at pH 5.0. The concentration of 0.6% (w/v) CNF and ChNF was chosen based on previous research (42). All sample mixtures (0.5 mL) were incubated for 24 h at room temperature prior to transferring 70 μL to a smooth 8 mm plate with 1 mm measuring gap for rheometry measurements using an Anton Paar Physica MCR 302 rheometer (Anton Paar, Graz, Austria). Temperature (23°C) was controlled with a Peltier hood H-PTD 200 (Anton Paar). To counter evaporation effects, the Peltier hood and a filled water ring were attached. The oscillatory tests consisted of three successive intervals: time sweep, frequency sweep and amplitude sweep. Time sweeps were performed at constant strain amplitude of 1% and angular frequency 1 s^-1^, frequency sweeps at angular frequencies from 100 to 1 s^-1^ (at constant strain amplitude 1%), and amplitude sweeps were performed in the strain amplitude range from 0.1 to 100% (at constant angular frequency 1 s^-1^). The time and frequency sweeps were performed at the linear viscoelastic region. The mechanical spectra for storage modulus (G’) and loss modulus (G’’) were recorded to determine the viscoelastic properties of the CNF and ChNF dispersions.

### 2.12 X-ray photoelectron spectroscopy (XPS)

XPS measurements were performed using chitin nanofibers (ChNF) suspended in 5 mM sodium acetate buffer (pH 3.5) to a final concentration of 0.6 % w/v; the ChNF was treated with equimolar amounts of either PcaLOOL2 (1 mg/mL) or PcaLOOL7 (0.5 mg/mL). All sample mixtures (0.5 mL) were incubated for 24 h at room temperature and freeze-dried prior to analysis. XPS analysis was performed using a Kratos AXIS Ultra DLD X-ray photoelectron spectrometer equipped with a monochromated AlKα X-ray source (1486.7 eV) run at 100 W. A pass energy of 80 eV and a step size of 1.0 eV were used for the survey spectra, while a pass energy of 20 eV and a step size of 0.1 eV were used for the high-resolution spectra. Photoelectrons were collected at a 90° take-off angle under ultra-high vacuum conditions, with a base pressure typically below 1 × 10-9 Torr. The diameter of the beam spot from the X-ray was 1 mm, and the area of analysis for these measurements was 300 µm x 700 µm. Both survey and high-resolution spectra were collected from three different spots on each sample surface in order to check for homogeneity and surface charge effects. All spectra were charge-corrected relative to the position of C–O bonding of carbon at 286.5 eV.

### 2.13 Assay for synergism in enzymatic polysaccharide hydrolysis

Synergism studies were performed in 2 mL Eppendorf tubes in a total reaction volume of 1 mL. Tested cellulose substrates (Avicel® PH-101 or Whatman® qualitative filter paper grade 1 disks created using a hole punch) were suspended in 50 mM sodium acetate buffer (pH 5.0) to a final concentration of 25 mg/ mL. Substrate suspensions were then treated in two ways: 1) with a mixture of a given PcaLOOL (0.041 mg/ mL) and the commercial cellulase Cellic®CTec2 (0.405 mg/ mL) for 24 h in a ThermoMixer®C set at 40°C and 1000 rpm, or 2) pretreated with a given PcaLOOL (0.041 mg/ mL) for 1 h in a ThermoMixer®C set at 25°C and 1000 rpm prior to addition of Cellic®CTec2 (0.405 mg/ mL), after which the incubation proceeded for 24 h at 40°C and 1000 rpm. BSA was used as a reference. Aliquots were sampled at regular time intervals and analysed by the para-hydroxybenzoic acid hydrazide (PAHBAH) assay. Substrate suspensions of Whatman® qualitative filter paper grade 1 were treated in three additional ways 1) with a mixture of a given PcaLOOL (0.7 mg/ mL) and the commercial cellulase Celluclast® (7 mg/ mL) for 24 h in a ThermoMixer®C set at 50°C and 1000 rpm, 2) with a mixture of a given PcaLOOL (0.05 mg/ mL) and a commercial *endo*-1,4-β-D-glucanase (0.5 mg/ mL) for 24 h in a ThermoMixer®C set at 50°C and 1000 rpm, or 3) pretreated with a given PcaLOOL (0.041 mg/ mL) for 72 h in a ThermoMixer®C set at 25°C and 1000 rpm prior to addition of Cellic®CTec2 (0.405 mg/ mL), after which the incubation proceeded for 2 h at 40°C and 1000 rpm. Aliquots were sampled at the end of the incubation period and analysed by the 3,5-dinitrosalicylic acid (DNS) assay calibrated against glucose (43).

Synergism experiments were also performed using chitin from shrimp shells and a commercial chitinase from *T. viride*. Chitin was suspended in 50 mM sodium acetate buffer (pH 6.0) to a final concentration of 25 mg/ mL and treated in two ways: 1) with a mixture of a given PcaLOOL (0.008 mg/ mL) and chitinase (0.08 mg/ mL) for 24 h in a ThermoMixer®C set at 25°C and 1000 rpm or 2) pretreated with a given PcaLOOL (0.75 mg/ mL) for 24 h in a ThermoMixer®C set at 25°C and 1000 rpm prior to addition of chitinase (0.02 mg/ mL), after which the incubation proceeded for another 24 h at the same conditions. BSA was used as a reference, aliquots sampled after 24 h were analyzed by the 3,5-dinitrosalicylic acid (DNS) assay calibrated against glucose.

### 2.14 Mycelia growth experiments

*Ganoderma lucidum* (reishi mushroom, HAMBI FBCC665) and *Pleurotus ostreatus* (oyster mushroom, HAMBI FBCC0515) were obtained from the HAMBI Culture Collection (University of Helsinki, Faculty of Agriculture and Forestry, Department of Microbiology). The isolated fungal cultures was previously identified with internal transcribed spacer polymerase chain reaction (ITS-PCR) (44). Fungal cultures were propagated and maintained on 2% (w/w) malt extract (LabM, Heywood, UK) in 2% (w/w) agar (Scharlab, Sentmenat, Spain) at 4°C ± 1°C. For mycelial growth experiments, 2 mL malt extract (2% w/w) in 4 mL screw neck vials (Fisher Scientific, Loughborough, UK) were inoculated with an agar piece of the *G. lucidum* or *P. ostreatus* culture. Malt extract was then supplemented with 0.1 mg/ mL PcaLOOL2, PcaLOOL7, or PcaLOOL12 prepared in 50 mM sodium acetate buffer (pH 5.0). Vials were sealed with cotton filters and open top screw caps (La-Pha-Pak, Langerwehe, Germany) and incubated at 21°C (± 1°C). Fungal growth was measured over 10 days at 25°C by multiple light scattering using a Turbiscan™ LAB and adapter for a 4 mL cell (Formulaction, Toulouse, France). Transmittance at 880 nm was recorded by a synchronous optical sensor that moves along the vertical axis at 40 µm intervals along the cylindrical measurement cell (45).

## 3. Results and discussion

### 3.1 Sequence analysis

Previous transcriptomic analysis of *P. carnosa* grown on aspen or spruce uncovered the expression of 12 loosenin-like proteins (LOOL), which were classified into phylogenetic subgroup A and subgroup B (28, 29). Whereas transcript abundancies were comparatively high for PcaLOOL12 during *P. carnosa* cultivation on wood, PcaLOOL2 transcript abundancies were highest overall (28, 29). PcaLOOL2 and PcaLOOL12 belong to subgroup B and are 25% and 34% identical to LOOS1 from *B. adusta*, respectively (28). In addition to selecting PcaLOOL2 and PcaLOOL12 for recombinant expression and characterization, PcaLOOL7 and PcaLOOL9 were selected from subgroup A, and are 64% and 59% identical to LOOS1, respectively.

The microbial expansin BsEXLX1 comprises a polysaccharide binding surface (PBS) lined with aromatic and polar residues that span the D1 and D2 domains of the protein (15, 46). Several of the D1 residues that form the PBS in BsEXLX1 are conserved in loosenins; including Gly38, Ala39, Ala52 and Asp93 of LOOS1 (Fig. 1; numbering of amino acids includes the N-terminal secretion signal). Notably, PcaLOOL2 and PcaLOOL12, as well as the N2 loosenin, harbour an insertion upstream of the PBS. PcaLOOL2 and PcaLOOL12 are further distinguished by an 8 – 9 amino acid deletion between residues equivalent to Gly63 and Pro72 in LOOS1.

**Figure 1:**
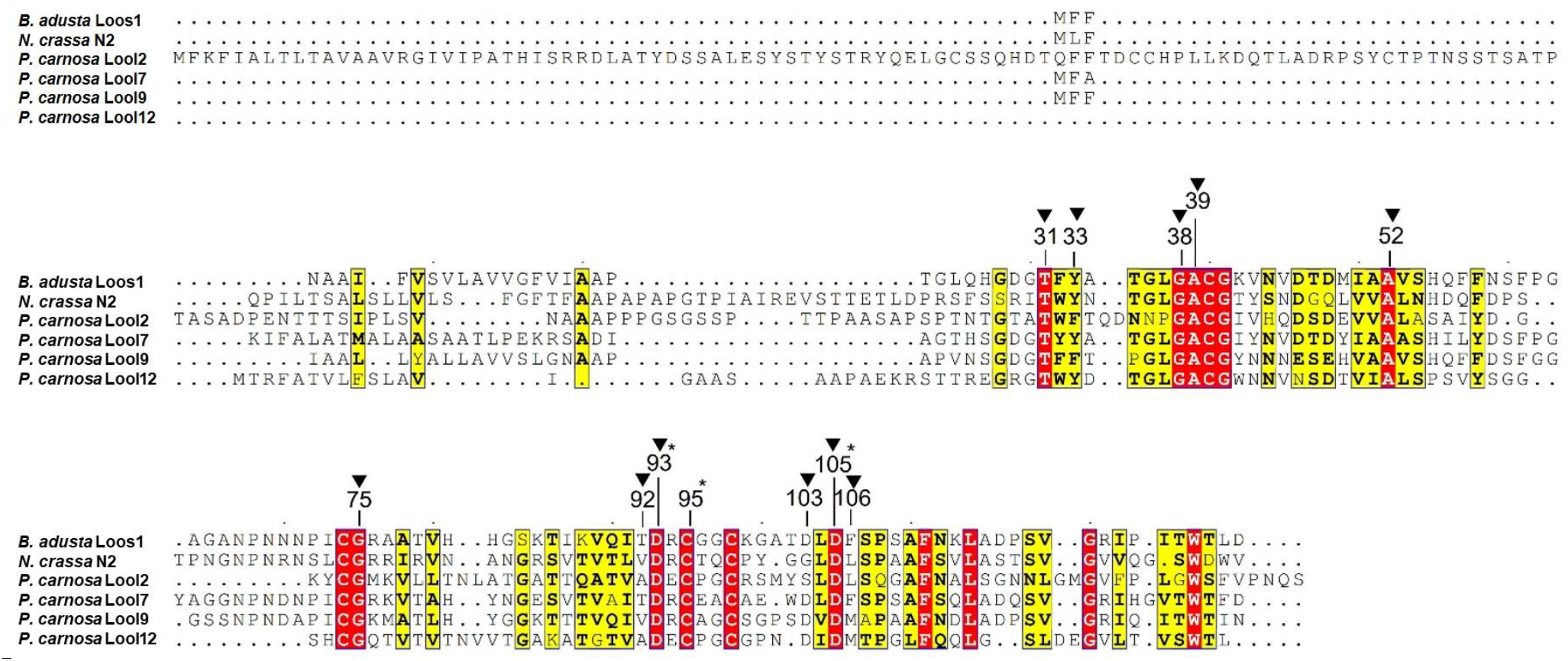
Multiple amino acid sequence alignment of *P. carnosa* LOOLs with characterized loosenins *B. adusta* LOOS1 and *N. crassa* N2. Strictly conserved regions are shown in red blocks, similar residues in yellow blocks. Grey boxes indicate chemical similarity across a group of residues. Numbering of amino acid residues corresponds to positions in LOOS1 and includeds the predicted N-terminal signal sequence. Triangles (▼) indicate residues contributing to form a shallow binding groove or hydrogen bonds. Stars (*) indicate residues proposed to be crucial for the activity of EXLX1. Amino acid sequences of *P. carnosa* PcaLOOL2 (GenBank code EKM55357.1), PcaLOOL7 (GenBank code EKM53490.1), PcaLOOL9 (GenBank code EKM52742.1), PcaLOOL12 (GenBank code EKM51974.1); *N. crassa* N2 (GenBank code XP_959591.1) and *B. adusta* Loos1 (GenBank code ADI72050.2) were retrieved from the NCBI protein database. Alignments were performed with Clustal Omega and the figure was generated with ESPript 3.

Thr12 and Asp82 in BsEXLX1 (previously reported numbering; does not include the N-terminal secretion signal) are predicted to form hydrogen bonds to glucan molecules and are strictly conserved among all microbial expansin-related proteins, including the selected PcaLOOLs (Fig. 1). An alanine substitution of Asp82 resulted in a complete loss of BsEXLX1 activity, whereas alanine substitution of Thr12 reduced BsEXLX1 activity by 70% (17); accordingly, the Asp82-Thr12 pair constitutes the presumptive active site of the D1 domain (12, 24). Asp82 in BsEXLX1 corresponds to the sole catalytic residue of GH105 lytic transglycosylases like EcMltA and to the proton donor in the catalytic site of GH45 enzymes. Thr12 in BsEXLX1 is equivalent to the Thr residue in the TWY motif of EXPA and the TFY motif of EXPB plant expansins. The hydroxyl group of Thr12 participates in a conserved hydrogen bond with the carboxylic group of Asp82 and is conserved in the catalytic site of GH45 enzymes and possibly MltA enzymes, where it is likely important for the proper positioning of the catalytic Asp (9, 15).

Notable differences between loosenins (and loosenin-like proteins) and the D1 domain of BsEXLX1 include substitution of Tyr97 in BsEXLX1 to a cysteine (e.g., Cys95 in LOOS1). Tyr97 in BsEXLX1 is important, but not essential, to cell wall creep (17).

Three other Cys residues are conserved among loosenins and sequence analyses with DISULFIND predict putative disulphide bond formation between positions 74 - 98 and 40 - 95. The N-terminal extension of PcaLOOL2 harbours four additional Cys residues that could form two additional disulphide bonds. While disulphide bridges are absent from BsEXLX1, plant expansins possess three disulphide bridges in D1, whose six participating Cys residues are well conserved among EXPA and EXPB families (9, 15, 16). ScEXLX1 from *Schizophyllum commune* also contains two disulphide bridges in D1 and the involved residues align with Cys40, Cys74, Cys95 and Cys98 of loosenins (47) (Suppl. Fig. S1).

### 3.2 Production of recombinant PcaLOOLs

Codon optimized genes encoding PcaLOOL2, PcaLOOL7, PcaLOOL9 and PcaLOOL12 were obtained as subcloned in pPICZαA and heterologously expressed in *P. pastoris* SMD1168H under control of a methanol-inducible promoter. The recombinant PcaLOOLs were secreted and purified by Ni-NTA affinity chromatography; the yield of PcaLOOL2, PcaLOOL7, PcaLOOL9 and PcaLOOL12 were 22 mg/ L, 28 mg/ L, 9 mg/ L and 27 mg/ L, respectively. The electrophoretic molecular weight of PcaLOOL2 and PcaLOOL9 were higher than expected, which was attributed to glycosylation in both cases, and potential formation of a PcaLOOL2 dimer (Suppl. Fig S2A). The identity of all protein bands was confirmed by MALDI-TOF MS analysis (Suppl. Fig. S2B, C). Moreover, circular dichroism (CD) spectroscopy indicated that all PcaLOOLs were well folded (Suppl. Fig. S3A). Briefly, all PcaLOOLs showed similar CD spectra indicating β-sheet-rich structures, consistent with the predicted DPBB fold (Suppl. Fig. S3B). The CD spectra of PcaLOOL2 and PcaLOOL7 also showed a positive peak at ∼220 nm, which has been assigned to the poly-L-proline type II (polyproline-II or PPII) type of secondary structure (48). During thermal unfolding, all PcaLOOLs underwent a single thermal transition as a function of temperature.

### 3.3. Lytic activity

The loosenins produced herein were tested for lytic activity, including hydrolytic activity. Consistent with previous analyses of expansin-related proteins, none of the PcaLOOLs characterized herein produced hydrolysis or lytic products from any of the tested polysaccharides, including xylan, xyloglucan, glucomannan, β-glucan, peptidoglycan, carboxymethyl cellulose, cellohexaose, and hexacetyl-chitohexaose (Suppl Fig. S4). The absence of hydrolytic activity in expansins has been attributed to the absence of a key aspartic acid residue (Asp10 in HiCel45A) that functions as the general base in many GH45 endoglucanases. In contrast, the catalytic proton donor (Asp121 in HiCel45A) is conserved in expansins and expansin-related proteins (Asp82 in BsEXLX1; Asp105 in LOOS1) (15, 17, 49). Molecular dynamics simulations of the HiCel45A D10N mutant indicated the possibility of a non-hydrolytic reaction mechanism when the catalytic base aspartic acid is missing, as it is the case in EcMtlA (50). EcMltA utilizes only one acidic amino acid to cleave the β-1,4-glycosidic bonds in bacterial cell wall peptidoglycan (51). Nevertheless, the apparent lack of lytic activity for PcaLOOLs is in line with extensive experimental evidence debunking the proposed lytic action of expansins (52).

### 3.4 Activity of PcaLOOLs on cellulose substrates

#### 3.4.1 Binding to cellulose

The binding capacities of the recombinant PcaLOOLs to cellulose were measured as a first step to evaluating their potential non-lytic action on cellulosic materials (Fig. 2A). Highest levels of binding to cellulose were measured for PcaLOOL2 at pH 5.0 to pH 6.0, where between 20-25% of PcaLOOL2 adsorbed to the cellulose substrate (Avicel® PH-101) after 1 h at room temperature. Similarly, Quiroz-Castañeda et al. (2011) (19) reported preferential adsorption of LOOS1 to Avicel® PH-101 in comparison to BSA. For all other PcaLOOLs, however, the extent of binding to Avicel PH101 was negligible and indistinguishable from binding values obtained using BSA. Similarly, the D1 domain of BsEXLX1 does not bind to Avicel (17).

**Figure 2:**
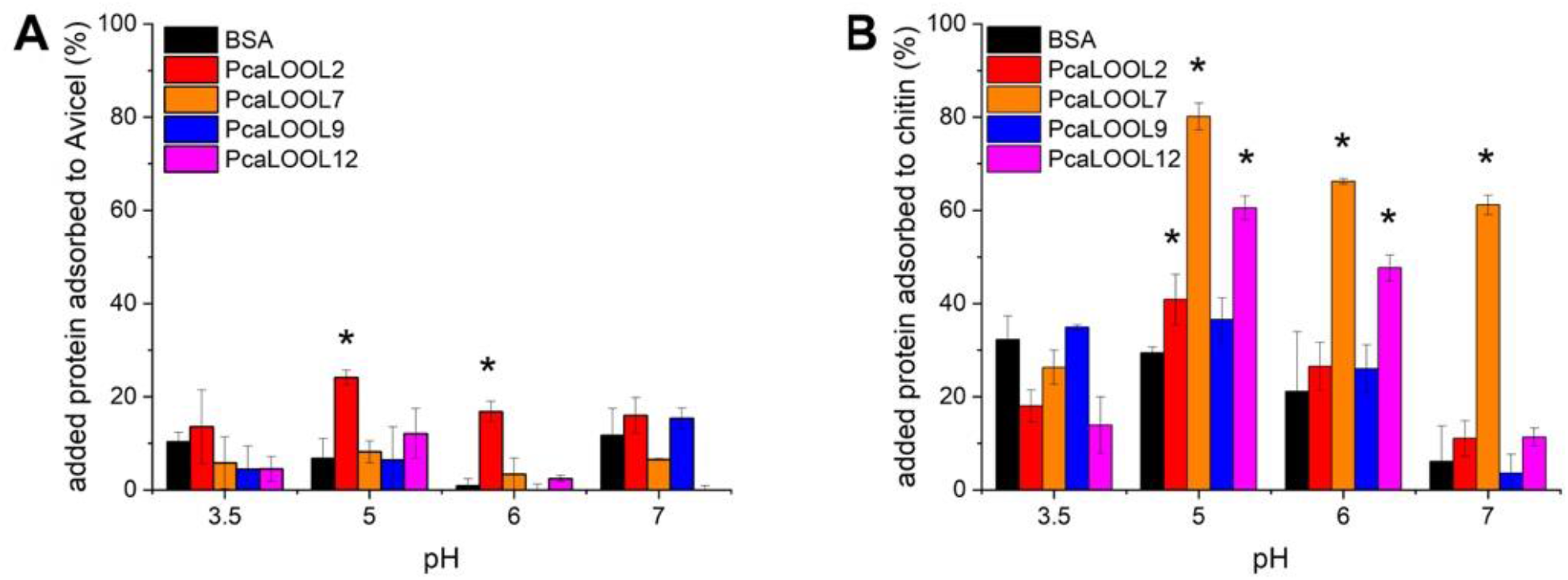
Adsorption of PcaLOOLs to cellulose and chitin. Affinity purified PcaLOOLs [0.1 mg/ mL] were incubated with 25 mg/ mL substrate for 1 h at room temperature in a tube rotator set at 20 rpm. Unbound protein in the supernatant was determined by the BCA assay. Percentage of added protein adsorbed to substrate was calculated considering protein and substrate blanks. BSA was used as a reference. Asterisks (*) indicate statistical significance (p ≤ 0.05; two-tailed T-test) of increased PcaLOOL adsorption compared to BSA. (A) Adsorption of PcaLOOLs to Avicel® PH-101 at 3.5, 5.0, 6.0 or 7.0; n = 3, errors correspond to standard deviation of the mean. (B) Adsorption of PcaLOOLs to chitin from shrimp shells at pH 3.5, 5.0, 6.0 or 7.0; n = 3, errors correspond to standard deviation of the mean.

#### 3.4.2 Weakening of filter paper

All four PcaLOOLs weakened cellulose filter paper as measured using the breaking force assay (53-55) (Fig. 3). In this assay, filter paper strips are incubated in buffered solutions containing the target protein and then clamped in an extensometer that records the force required to break the material (41). Compared to the buffer treatments, the application of PcaLOOL7, PcaLOOL9 and PcaLOOL12 significantly (p ≤ 0.05) decreased the breaking force of filter paper strips by 11-16%, compared to a 30% reduction with BsEXLX1 (Fig. 3). The low binding to Avicel and yet ability to weaken cellulose filter paper suggests ability to desorb from cellulose surfaces is important to PcaLOOL function. Notably, previous mutagenesis of the D2 domain of BsEXLX1 revealed a non-linear relation between microbial expansin binding to cellulose and filter paper weakening (56).

**Figure 3:**
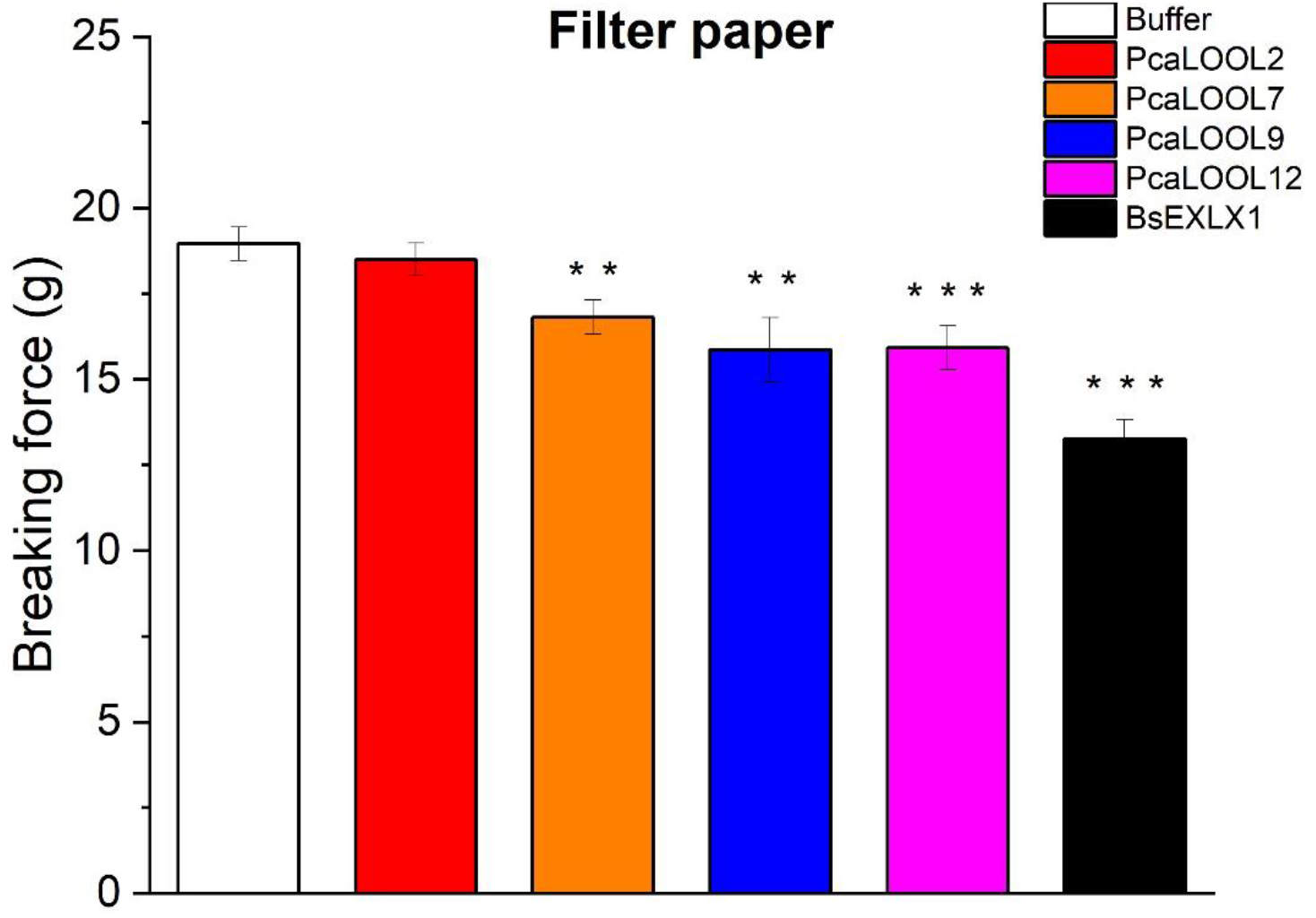
Breaking strength of filter paper after treatment with PcaLOOLs. Strips of Whatman® qualitative filter paper grade 3 were treated with or without 0.2 mg/ mL protein in 20 mM MES buffer, pH 6, for 4 h at 25°C. Strips were extended at 1.5 mm/min and the breaking force recorded. BsEXLX1 was used as a positive reference. Error bars are +/-standard error of the mean, n =34 for buffer control; 25 for BsEXLX1 and 15-19 for different PcaLOOL proteins. Asterisks (** or ***) indicate statistical significance (p < 1% or p < 0.01%, respectively).

#### 3.4.3 Impact of PcaLOOLs on cell wall creep

The defining characteristic of plant expansins and microbial expansin-like proteins is their ability to induce time-dependent, irreversible elongation (creep) of plant cell walls. The corresponding plant cell wall creep assay is the most diagnostic, specific, and sensitive method to detect expansin activity (12, 17). While the induction of cell wall creep correlates well with weakening of filter paper in BsEXLX1 and its mutants, this connection is much weaker for other microbial expansin-related proteins (17, 53). Indeed, none of the PcaLOOLs investigated herein were able to increase the extension rate of wheat coleoptile cell walls, a finding in accordance with the behavior of other single-domain DPBB proteins (10) (Suppl. Fig. S5).

#### 3.4.4 Impact of PcaLOOLs on the visco-elastic behaviour of cellulose nanofibers (CNF)

Rheology measurements were obtained to further quantify the impact of PcaLOOLs on cellulose fiber weakening. First, the linear viscoelastic region (LVE) was determined, after which, time sweep, frequency sweep, and amplitude sweep measurements were performed. For all experiments using CNF, the storage (G’) and loss (G’’) moduli did not change significantly over time (Suppl. File. S1). The expected gel-like behavior of the starting CNF was further confirmed by the frequency sweep where G’ was invariably higher than G’’ (Suppl. File S2; 57) and the amplitude sweep where G’ dropped below G’’ at high strain (i.e., at the limiting value of the LVE region) (Suppl. File S3). The limiting value of the LVE specifies the yield stress of the material, which is the peak value of the elastic stress (σ’), calculated as σ’= G’γ, where G’ is the storage modulus and γ is the strain amplitude (58, 59). The corresponding strain amplitude at the yield stress is defined as the yield strain (Suppl. Files. S4 and S5).

All PcaLOOLs lowered the yield strain of the CNF beyond negligible impacts observed using BSA (Fig. 4 A and B), consistent with LOOL-mediated disruption of the fibrillar network. At pH 5.0, the impact of PcaLOOLs on the CNF network increased with increasing protein concentration (Fig. 4A). A decrease in yield strain of CNF was also observed after treatment with PcaLOOLs at pH 3.5; however, a clear correlation between yield strain and protein concentration was not observed at that pH value.Notably, at pH 3.5 and comparatively low protein concentrations, the BSA treatment increased the yield strain of CNF (Fig. 4B), which can be explained by the expected unfolding and resulting increase in surface charge previously reported for BSA at acidic conditions (60).

**Figure 4:**
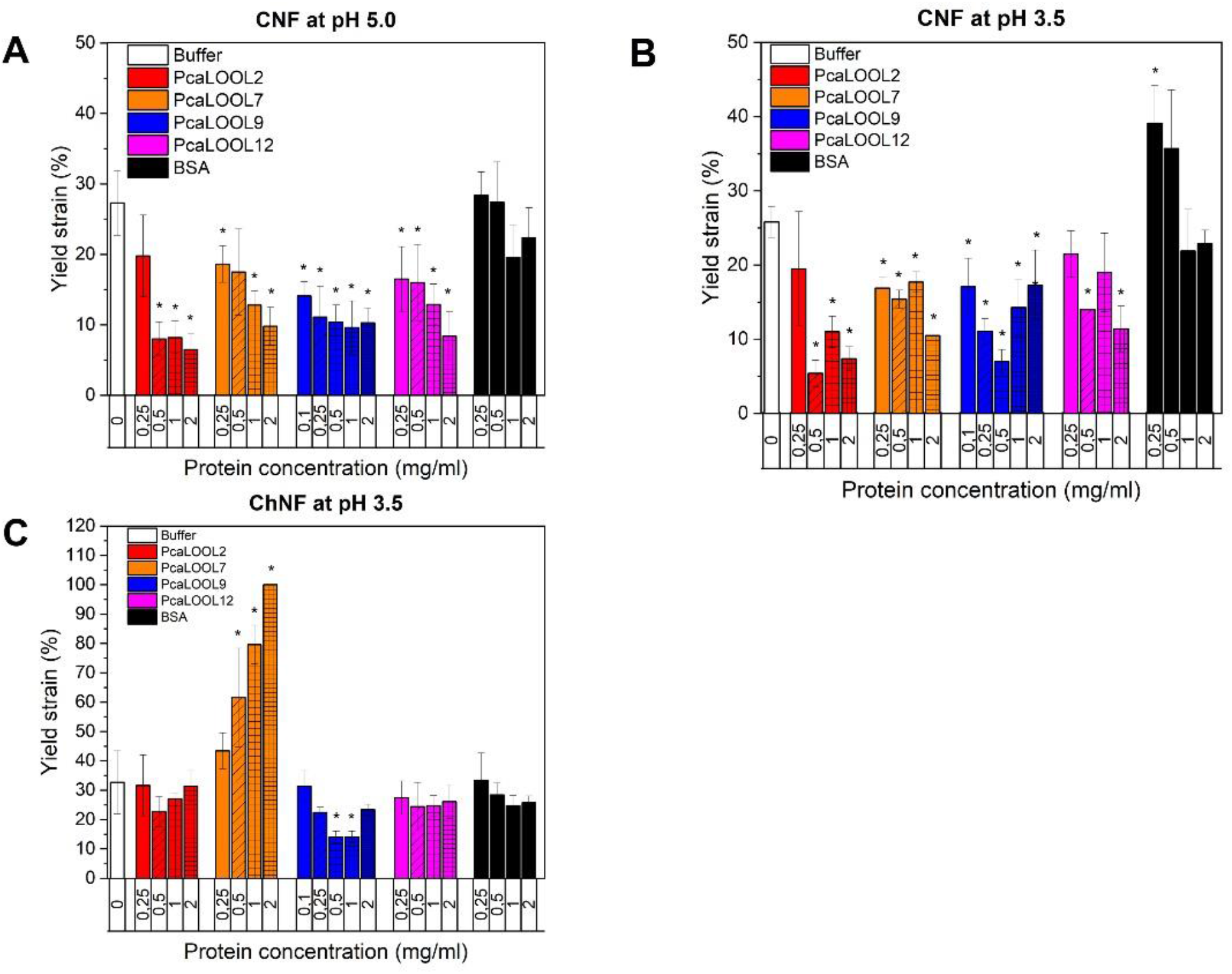
Yield strain determination, measured by means of oscillatory rheological tests. 0.6% substrate was treated with increasing concentrations of protein for 24 h at room temperature and subjected to rheological tests. (A) CNF treated with PcaLOOLs at pH 5.0 (B) CNF treated with PcaLOOLs at pH 3.5. (C) ChNF treated with PcaLOOLs at pH 3.5. Asterisks (*) indicate statistical significance (p ≤ 0.05; two-tailed T-test) of decreased yield strain upon PcaLOOL treatment of CNF compared to only buffer. n = 2-5, errors correspond to standard deviation.

Considering the pKa of Asp is 3.9, the putative ‘catalytic’ Asp residue of PcaLOOLs should be largely protonated at pH 3.5 and deprotonated at pH 5.0. Accordingly, the current results suggest PcaLOOLs function better at pH values where the catalytic Asp is deprotonated. Similarly, cell wall creep by BsEXLX1 was highest above pH 5.5 (until pH 9.5, where the protein begins to precipitate) (17).

#### 3.4.5. Impact of PcaLOOLs on cellulase action

Each of the PcaLOOLs produced herein was tested for ability to boost Cellic®CTec2, Celluclast® or *endo*-1,4-β-D-glucanase action on Avicel® PH-101 and Whatman® qualitative filter paper. Notably, despite their ability to weaken filter paper and CNF networks, none of the PcaLOOLs substantially increased cellulase action on selected cellulosic substrates (Suppl. Fig. S6, S7). Indeed, reported impacts on biomass deconstruction vary considerably and depend strongly on the biomass source (26, 27, 61). For example, previous experiments showing complementary action between LOOS1 and a commercial cellulase were performed using cotton fiber and *Agave tequilana* fibers (19). Since PcaLOOL2 and PcaLOOL12 were recently shown to boost xylanase migration through cellulose-xylan composites (62), future studies of PcaLOOL impacts on the enzymatic deconstruction of cellulosic substrates should include complex lignocellulosic materials from diverse biomass sources.

### 3.5 Activity of PcaLOOLs on chitin substrates

#### 3.5.1 Binding to chitin

Previous co-expression analyses clustered PcaLOOL2 with a predicted carbohydrate esterase family 9 N-acetyl-glucosamine 6-phosphate deacetylase, PcaLOOL7 with a putative glycoside hydrolase family 18 chitinase, and PcaLOOL12 with a predicted glycosyl transferase family 2 chitin synthase (29). These earlier observations point to a possible role for PcaLOOLs in fungal cell wall morphogenesis, potentially through targeting chitin in fungal cell walls during hyphal growth. A clear pH dependency was observed for PcaLOOL binding to chitin, where after 1 h at room temperature highest binding in all cases was observed between pH 5.0 and 6.0 (Fig. 2B). Despite the clear binding to chitin, and highest binding by PcaLOOL7 followed by PcaLOOL12, none of the PcaLOOLs produced herein subtantially boosted chitinase action on chitin substrates (Suppl. Fig. S8).

#### 3.5.2 Impact of PcaLOOLs on the visco-elastic behaviour of chitin nanofibers (ChNF)

Similar to the rheological response of CNF, the ChNF samples were viscoelastic solids with a typical gel-like behavior (Suppl. File S1 and S2). Accordingly, rheology measurements were obtained using ChNF to further quantify the impact of PcaLOOLs on chitin fibrillar networks. In this case, the rheology measurements could only be performed at pH 3.5 so as to retain the expected degree of acetylation within the ChNF substrate. At pH 3.5, binding to chitin was highest for PcaLOOL9 (Fig. 2B); PcaLOOL9 also induced the greatest loss in yield strain of the ChNF network (Fig. 4C, Suppl. File S6). Strikingly, PcaLOOL7 increased the yield strain of ChNF in a dose dependent manner. Since PcaLOOL7 behaved similarly to other PcaLOOLs on CNF at pH 3.5, the impact of PcaLOOL7 on ChNF appears to be substrate rather than pH dependent. It is conceivable that rather than disrupting interfibrillar associations, PcaLOOL7 disrupts irregularities (e.g., intrafibrillar dislocations) within ChNF or interacts with two neighbouring ChNF fibrils, leading to more connection points within the ChNF network.

To further evaluate the impact of PcaLOOL7 on ChNF, the chemical state of ChNF surface elements was analysed by XPS before and after treatment with PcaLOOL7. For comparison with a PcaLOOL from a different subfamily, XPS analyses also included ChNF samples treated with equimolar concentrations of PcaLOOL2. Survey spectra of untreated ChNF confirmed the expected carbon, oxygen and nitrogen contents (i.e., ∼ 60 % carbon and 35-40 % oxygen and nitrogen combined (data not shown). The deconvolution of C 1s signals identified 3 peaks corresponding to C–C/C–H bonding (at 285 eV), C–N bonding (at 286 eV) and O–C–O (at 288.7 eV) (63, 64). The C 1s spectra of PcaLOOL2-treated ChNF revealed a significant increase in relative signal from C–N bonding and C–C/C–H bonding (Suppl. Fig. S9B), indicative of increased exposure of hydrophobic regions at the ChNF surface (65). This impact of PcaLOOL treatment of ChNF was much less pronounced for PcaLOOL7 (Suppl. Fig. S9C), consistent with the reorganization but reduced structural breakdown of the ChNF network. To further evaluate the differing impacts of PcaLOOL on chitin networks, their potential to modify complex biological samples was explored.

#### 3.5.3 Impact of PcaLOOLs on fungal growth

The cell walls of most filamentous fungi comprise interwoven microfibrils of chitin and β-1,3-glucans that are embedded in α-1,3-glucans and glycoproteins (66). Accordingly, the comparatively high binding of PcaLOOLs to chitin and varying influence of PcaLOOLs on the ChNF network motivated preliminary experiments that investigate impacts of PcaLOOLs on mycelial growth. Since there is no readily available genetic system for manipulating *P. carnosa*, the impact of the PcaLOOLs analyzed herein on the phenotype of filamentous fungi was assessed by external protein supplementation to fungal cultivations.

Like *P. carnosa, Ganoderma lucidum* is a white rot basidiomycete belonging to the class of Agaricomycetes and secretes a similar set of enzymes. *G. lucidum* is also one of the most common fungal strains used for the production of mycelium composite materials (44, 67). BLAST searches of available *G. lucidum* sequences did not detect significant similarities to PcaLOOL7, PcaLOOL2 or PcaLOOL12, which reduced the possibility that native proteins would mask the impact of the supplemented PcaLOOLs. *G. lucidum* was thus grown in liquid malt extract supplemented with 0.1 mg/ mL protein and mycelia growth was followed over 10 days by light scattering (signal over sample height). In comparison to the transmission of *G. lucidum* reference cultures, the transmission of *G. lucidum* cultures amended with PcaLOOL7 and PcaLOOL12 increased between days 5 to 10 (Fig. 5A). The impact of PcaLOOLs on fungal growth was similar in cultivations of the white-rot *Pleurotus ostreatus* (Fig. 5B). Since no measureable difference in fungal rate was observed in *G. lucidum* and *P. ostreatus* cultivations amended with PcaLOOLs, it is conceivable that differences in the transmittance of cultures amended with PcaLOOL12, and to a lesser extent PcaLOOL7, results from changes to the density of cultivated mycelia.

**Figure 5:**
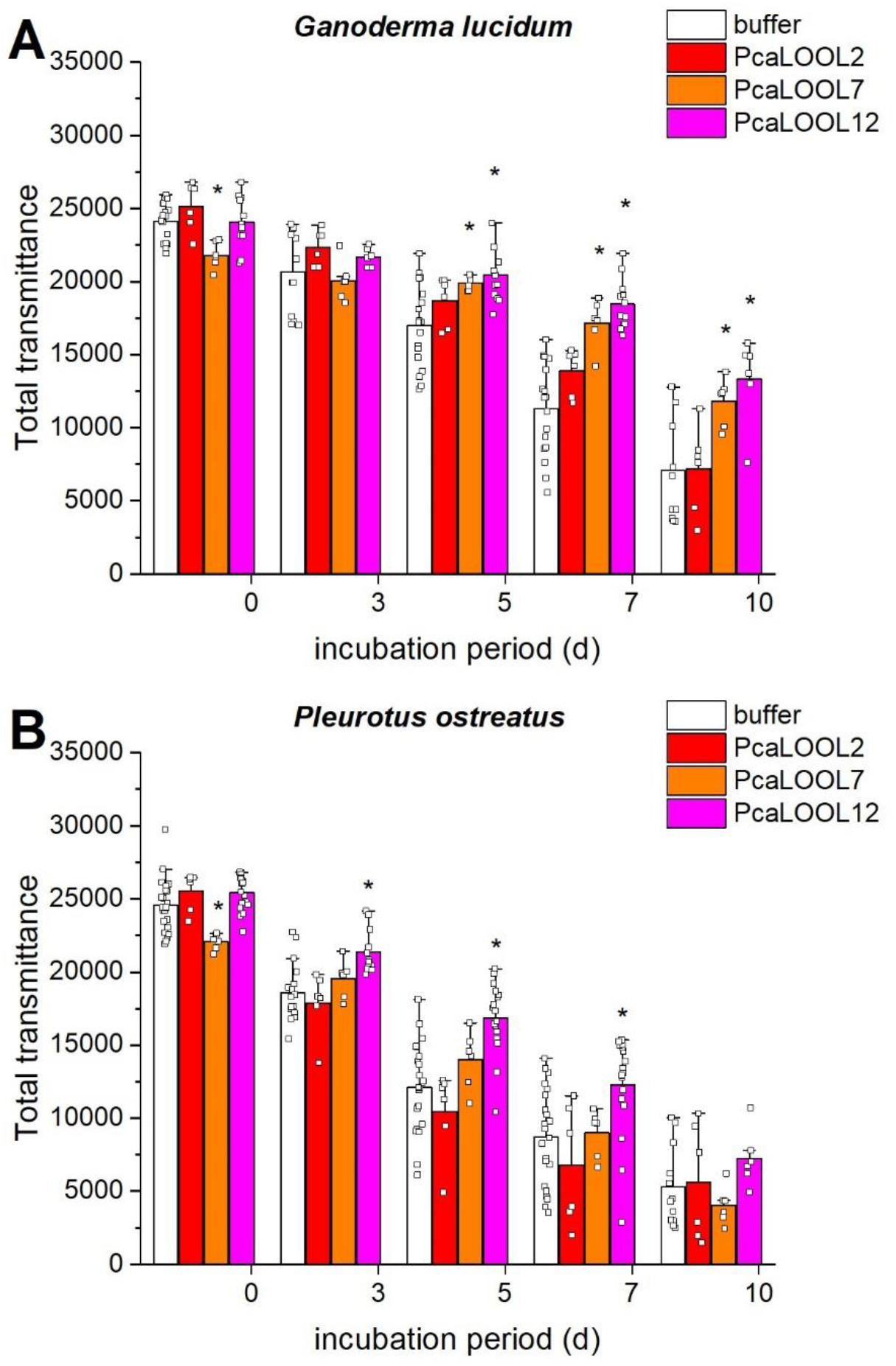
Effect of PcaLOOLs on the transmission of *G. lucidum* and *P. ostreatus* cultures. Malt liquid (2 mL) was inoculated with fungal mycelium and grown at 21°C for up to 10 days with 0.1 mg/ mL PcaLOOL. Fungal growth was monitored at 880 nm; n ≥ 6, errors correspond to standard deviation. Asterisks (*) indicate statistical significance (*p* ≤ 0.05; two-tailed t-test) of increased total transmission upon PcaLOOL supplementation compared to only buffer supplementation. (A) Total transmission of *G. lucidum* cultures containing PcaLOOL2 (red), PcaLOOL7 (orange), or PcaLOOL12 (magenta). (B) Total transmission of *P. ostreatus* cultures containing PcaLOOL2 (red), PcaLOOL7 (orange), or PcaLOOL12 (magenta).

## 4. Conclusions

The direct comparison of four loosenin-like proteins from a single filamentous fungus, *Phanerochaete carnosa*, uncovered functional characteristics that were common to all selected proteins and thereby deepen our understanding of microbial expansin-related proteins that comprise only the D1 (GH45-like) domain. In particular, all bound more strongly to alpha-chitin than to selected celluloses and none of the PcaLOOLs displayed lytic activity. Despite only weak binding to cellulose, all of the PcaLOOLs nevertheless reduced the yield strain of cellulose nanofibril networks. The lack of synergistic action between PcaLOOLs and cellulolytic enzymes observed herein likely reflects the comparative purity of the cellulose substrates used, and points to the importance of testing complex lignocellulosic substrates for application trials. The impact of PcaLOOLs on chitin fiber networks was more variable. For example, PcaLOOL7 increased the yield strain of ChNF whereas PcaLOOL9 had the opposite impact.

Preliminary studies to evaluate the impact of PcaLOOLs on fungal growth point to a possible biological function in fungal cell wall morphogenesis. This especially intriguing result provides new direction to fundamental studies that investigate the biological role of microbial expansin-related proteins, and opens new application concepts for their use in fungal biomass processing.

