## Supplemental Figures for "Loosenin-like proteins from *Phanerochaete carnosa* impact both cellulose and chitin fiber networks"

### SUPPORTING INFORMATION

**Mareike Monschein[a], Eleni Ioannou[a], Leamon AKM AL Amin[a], Jutta Varis[b], Edward R. Wagner[c], Kirsi S. Mikkonen[b], Daniel J. Cosgrove[c] and Emma R. Master[a,d,*]**

[a] Department of Bioproducts and Biosystems, Aalto University, Kemistintie 1, 02150 Espoo, Finland.
[b] Department of Food and Nutrition, University of Helsinki, Agnes Sjöbergin katu 2, 00014, Helsinki, Finland.
[c] Department of Biology and Center for Lignocellulose Structure and Formation, 208 Mueller Laboratory, Pennsylvania State University, University Park, State College, PA 16802, United States.
[d] Department of Chemical Engineering and Applied Chemistry, University of Toronto, 200 College Street, Toronto, Ontario, M5S 3E5, Canada.
*Corresponding author. phone number: +1 416-946-7861

**Supporting figure legends**

**Figure S1: Multiple amino acid sequence alignment of *P. carnosa* LOOLs with loosenins and non-catalytic module family EXPNs.** Strictly conserved regions are shown in red blocks, similar residues in yellow blocks. Grey boxes indicate chemical similarity across a group of residues. Numbering of amino acid residues corresponds to positions in LOOS1. Alignment includes *P. carnosa* PcaLOOL2 (GenBank code EKM55357.1), PcaLOOL7 (GenBank code EKM53490.1), PcaLOOL9 (GenBank code EKM52742.1), PcaLOOL12 (GenBank code EKM51974.1); *Neurospora crassa* N2 (GenBank code XP_959591.1); *Bjerkandera adusta* Loos1 (GenBank code ADI72050.2); *Mucor lusitanicus* EXPN (GenBank code KAF1805892.1); *Melampsora larici-populina* EXPN (GenBank code XP_007414134.1); *Schizophyllum commune* EXPN (GenBank code XP_003036918.1); *Sphaerosporella brunnea* EXPN (GenBank code KAA8896207.1); *Auriculariopsis ampla* EXPN (GenBank code TRM62543.1); *Endogone sp*. EXPN (GenBank code RUS20349.1); *Testicularia cyperi* EXPN (GenBank code PWZ03653.1); *Lentinula edodes* EXPN (GenBank code GAW08456.1); *Mycena chlorophos* EXPN (GenBank code GAT51218.1); *Phakopsora pachyrhizi* EXPN (GenBank code ALL40755.1); *Schizopora paradoxa* EXPN (GenBank code KLO11493.1); *Trametes versicolor* (GenBank code XP_008037594.1); *Heterobasidion irregulare* EXPN (GenBank code XP_009541212.1); *Fistulina hepatica* EXPN, partial (GenBank code KIY43622.1); *Moniliophthora roreri* EXPN (GenBank code ESK91981.1); *Mucor ambiguus* EXPN (GenBank code GAN00836.1); *Armillaria gallica* EXPN (GenBank code PBK82862.1); *Rickenella mellea* EXPN (GenBank code TDL27985.1); *Dendrothele bispora* EXPN (GenBank code THU87072.1); *Pyrrhoderma noxium* EXPN (GenBank code PAV22929.1). Corresponding amino acid sequences were retrieved from the NCBI protein database. Alignments were performed with Clustal Omega and the figure was generated with ESPript 3. Figure depicts detail of MSA covering conserved regions.

**Figure S2: Confirmation of PcaLOOL identity and purity.** (A) SDS-Page of recombinantly expressed *P. carnosa* LOOLs.  Affinity-purified proteins (20 µg) were separated on a 4 - 20% Mini-PROTEAN® TGX™ gel and stained with Coomassie G-250 using the PageBlue™ Protein Staining Solution. Lane 1: PageRuler™ Prestained Protein Ladder; lane 2: PcaLOOL2; lane 3: PcaLOOL7; lane 4: PcaLOOL9; lane 5: PcaLOOL12; lane 6: PageRuler™ Unstained Low Range Protein Ladder. (B) MALDI-TOF mass spectra of peptide mixtures obtained by chymotrypsin digest of PcaLOOLs. The X and Y-axis depict molecular masses in Dalton and peak intensities. Matched peptide peaks are marked according to their molecular masses. A. PcaLOOL2, B. PcaLOOL7, C. PcaLOOL9, D. PcaLOOL12. (C) Amino acid sequence coverage of PcaLOOLs by peptides identified from MALDI-TOF MS analysis. Bold red letters indicate amino acid sequence coverage by peptides detected from MALDI-TOF MS analysis of the chymotrypsin digested PcaLOOLs. Sequences analysed by tandem mass spectrometry (MS/MS) are underlined. A. PcaLOOL2, B. PcaLOOL7, C. PcaLOOL9, D. PcaLOOL12.

**Figure S3: Secondary structure analysis of PcaLOOLs.** (A) Far UV CD spectra. CD data of affinity purified PcaLOOLs [0.1 mg/ mL] were collected between 190 and 280 nm at 22°C using a 0.1 cm path-length quartz cuvette. Raw data were averaged, smoothed and the buffer baseline subtracted, using the Chirascan Pro-Data Viewer

(Applied Photophysics) software. Global 3 software was used for calculating thermal transitions. Mean residue molar ellipticity ([θ]MR) is shown at selected wavelengths; n = 3. (B) Estimated secondary structure content (%) calculated by the BeStSel web server and melting temperatures (Tm, °C) analyzed with Global3 using CD data.

**Figure S4: Evaluating the potential of PcaLOOLs to depolymerize polysaccharides.** (A) Test for hydrolytic activity. Xylan, CMC or glucomannan [0.5% (w/v)] were incubated with 0.01 mg/ mL protein in 50 mM sodium acetate buffer (pH 5.0) for 16 h at 40°C and 700 rpm. Reducing end concentrations were determined by the PAHBAH assay. Supplementation of buffer only was used as a reference. n=3, errors correspond to standard deviation of mean. Turbidity measures of lytic activity towards β-glucan (B) and peptidoglycan (C). Substrates [0.35 mg/ mL] were incubated with 0.05 mg/ mL protein in 50 mM sodium acetate buffer (pH 5.0) for 0 - 24 h at 50°C and 1000 rpm. Supplementation of buffer only or BSA was used as a reference. Absorbance of reactions measured at 600 nm, $n \geq 2$, errors correspond to standard deviation. (D) Thin layer chromatography (TLC) measures of lytic activity towards peptidoglycan, β-glucan, cellohexaose, chitohexaose or xyloglucan substrates. Supplementation of buffer only, BSA, *endo*-1,4-β-D-glucanase or hen egg-white lysozyme was used as a reference. Samples were collected after 24 h and spotted on TLC silica gels, developed in propanol/ 25% ammonia (2:1) and visualized with 10% sulfuric acid in ethanol. Lane 1: PcaLOOL2; lane 2: PcaLOOL7; lane 3: buffer; lane 4: PcaLOOL9; lane 5: PcaLOOL12; lane 6: BSA; lane S1: N-acetylglucosamine standard (peptidoglycan), glucose standard (β-glucan, xyloglucan), *endo*-1,4-β-D-glucanase (cellohexaose), hen egg-white lysozyme (chitohexaose); lane S2: xylose standard (xyloglucan).

**Figure S5: Test for wall extension activity.** Extension rates of alkali pretreated wheat coleoptile cell wall upon treatment with 0.2 mg/ mL PcaLOOLs in 20 mM MES buffer (pH 6.0), applied at the time indicated by arrows. Buffer without addition of protein was used as a negative control. Each curve represents the average rates of four coleoptile walls.

**Figure S6: Enzymatic hydrolysis of Whatman® filter paper upon supplementation of PcaLOOLs.** Supplementation of buffer only or BSA was used as a reference; reducing sugar concentrations were determined by the DNS assay. (A) Pre-incubation of Whatman® qualitative filter paper grade 1 [25 mg/ mL] with 0.041 mg/ mL PcaLOOLs at 25°C and 1000 rpm for 72 h, followed by enzymatic hydrolysis with 0.405 mg/ mL Cellic®CTec2 at 40°C, 1000 rpm for 2 h. n = 3. (B) Enzymatic hydrolysis of Whatman® qualitative filter paper grade 1 [25 mg/ mL] with 7 mg/ mL Celluclast®, supplemented with 0.7 mg/ mL LOOLs, at 50°C and 1000 rpm for 24 h. n = 3. (C) Enzymatic hydrolysis of Whatman® qualitative filter paper grade 1 [25 mg/ mL] with 0.5 mg/ mL *endo*-1,4-β-D-glucanase supplemented with 0.05 mg/ mL PcaLOOLs, at 50°C and 1000 rpm for 24 h. n = 3.

**Figure S7: Enzymatic hydrolysis of cellulosic substrates upon supplementation of PcaLOOLs**. Reducing sugar concentrations were determined by the PAHBAH assay. (A) Enzymatic hydrolysis of Avicel® PH-101 [25 mg/ mL] with 0.405 mg/ mL Cellic®CTec2, supplemented with 0.041 mg/ mL LOOLs, at 40°C, 1000 rpm for 24 h. Supplementation of buffer only or BSA was used as a reference; n ≥ 3. (B) Pre-incubation of Avicel® PH-101 [25 mg/ mL] with 0.041 mg/ mL PcaLOOLs at 25°C and 1000 rpm for 1 h, followed by enzymatic hydrolysis with 0.405 mg/ mL Cellic®CTec2 at 40°C, 1000 rpm for 24 h. Supplementation of buffer only or BSA was used as a reference; n ≥ 3. (C) Pre-incubation of Whatman® filter paper grade 1 [25 mg/ mL] with 0.041 mg/ mL PcaLOOLs at 25°C and 1000 rpm for 1 h, followed by enzymatic hydrolysis with 0.405 mg/ mLCellic®CTec2 at 40°C and 1000 rpm for 24 h. Supplementation of buffer only or BSA was used as a reference; n = 3.

**Figure S8: Enzymatic hydrolysis of chitin upon supplementation of PcaLOOLs.** Reducing sugar concentrations were determined by the DNS assay. (A) Enzymatic hydrolysis of chitin from shrimp shells [25 mg/ mL] with 0.08 mg/ mL chitinase from *T. viride*, supplemented with 0.008 mg/ mL PcaLOOLs, at 25°C and 1000 rpm for 24 h. Supplementation of buffer only or BSA was used as a reference; n = 3. (B) Pre-incubation of chitin from shrimp shells [25 mg/ mL] with 0.75 mg/ mL PcaLOOLs at 25°C and 1000 rpm for 24 h, followed by enzymatic hydrolysis with 0.02 mg/ mL chitinase from *T. viride* at 25°C and 1000 rpm for 24 h. Supplementation of buffer only or BSA was used as a reference; n = 3.

**Figure S9: High resolution X-ray photoelectron C 1s spectra deconvolution of ChNF after treatment with equimolar quantities of PcaLOOL2 and PcaLOOL7.** (A) 0.6% ChNF suspended in 5 mM sodium acetate buffer (pH 3.5). (B) 0.6% ChNF treated with PcaLOOL2 (1 mg/ mL) at pH 3.5. (C) 0.6% ChNF treated with PcaLOOL7 (0.5 mg/ mL) at pH 3.5. In all cases, incubations were for 24 h at room temperature. The contributions of C–C/C–H, C–N, and O–C–O peaks are colored in green, red, and blue, respectively. The relative intensities of the signals are shown as relative %. Their sum is colored in black.

### Supporting figures

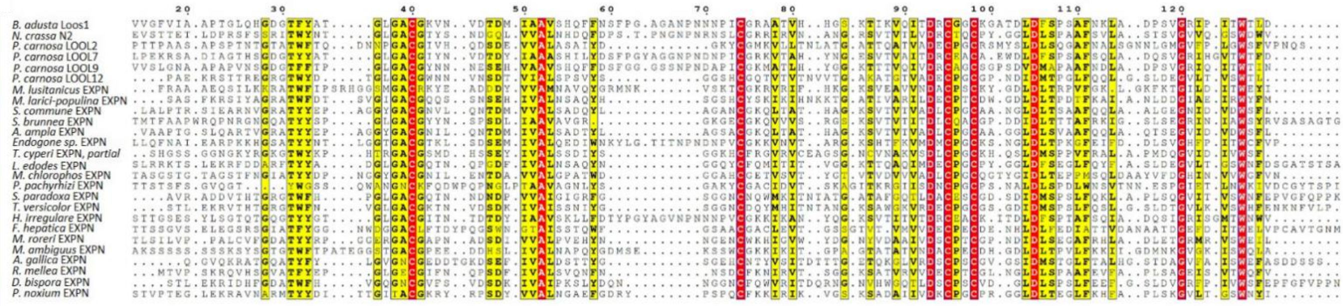


**Figure S1**

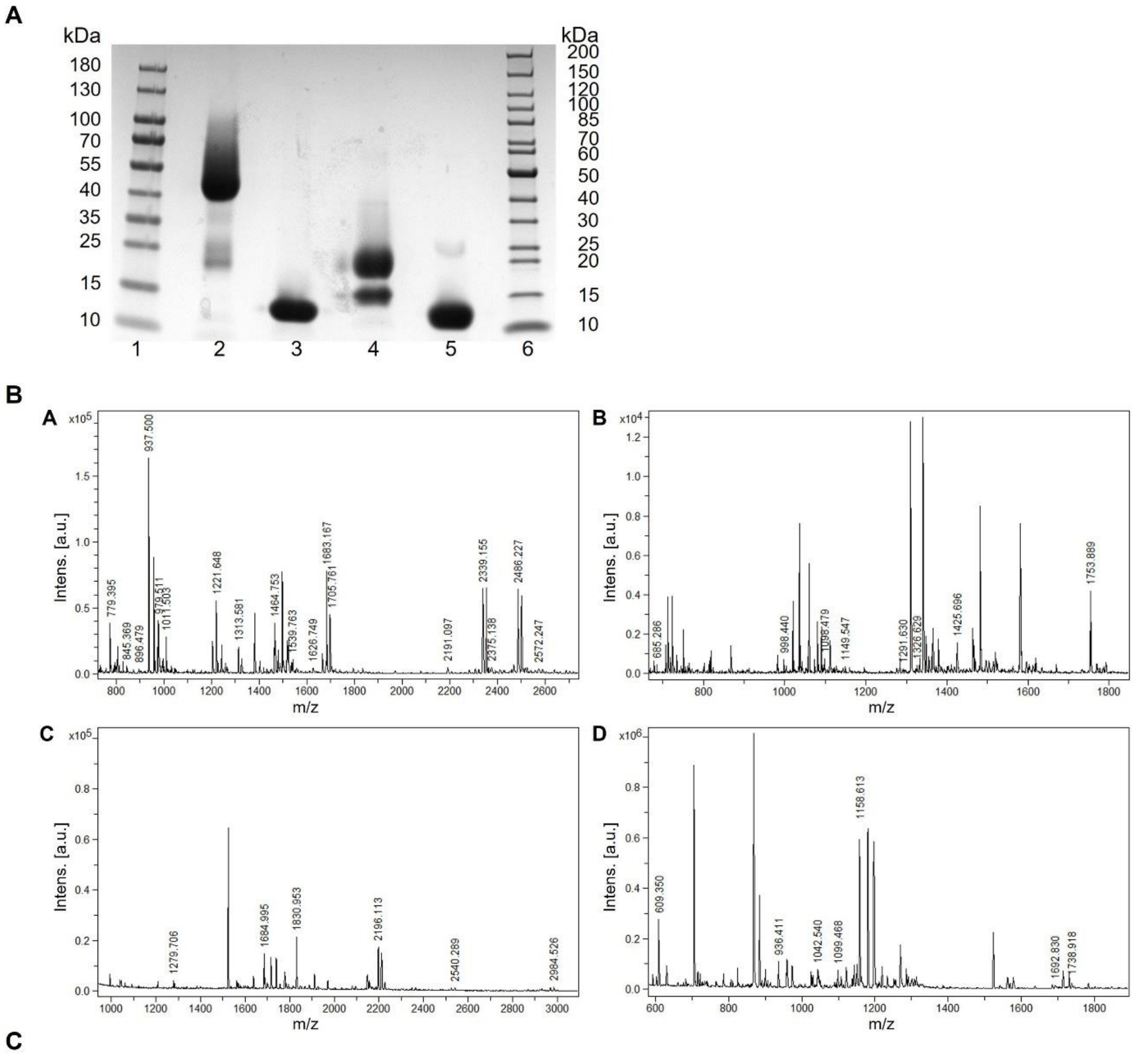


C

```
A IVIPATHISR RDLATYDSSA LESYSTYSTR YQELGCSSQH DTQFFTDCCH PLLKDQTLAD    60
  RPSYCTPTNS STSATPTASA DPENTTTSIP LSVNAAAPPP GSGSSPTTPA ASAPSPTNTG   120
  TATWFTQDNN PGACGIVHQD SDEVVALASA IYDGKYCGMK VLLTNLATGA TTQATVADEC   180
  PGCRSMYSLD LSQGAFNALS GNNLGMGVFP LGWSFVPNQS HHHHHH                  226

B ATLPEKRSAD IAGTHSGDGT YYATGLGACG IYNVDTDYIA AASHILYDSF PGYAGGNPND    60
  NPICGRKVTA HYNGESVTVA ITDRCEACAE WDLDFSPSAF SQLADQSVGR IHGVTWTFDH   120
  HHHHH                                                              125

C APAPVNSGDG TFFTPGLGAC GYNNNESEHV AAVSHQFFDS FGGGSSNPND APICGKMATL    60
  HYGGKTTTVQ IVDRCAGCSG PSDVDMAPAA FNDLADPSVG RIQITWTINH HHHHH        115

D APAEKRSTTR EGRGTWYDTG LGACGWNNVN SDTVIALSPS VYSGGSHCGQ TVTVTNVVTG    60
  AKATGTVADE CPGCGPNDID MTPGLFQQLG SLDEGVLTVS WTLHHHHHH              109
```

**Figure S2**

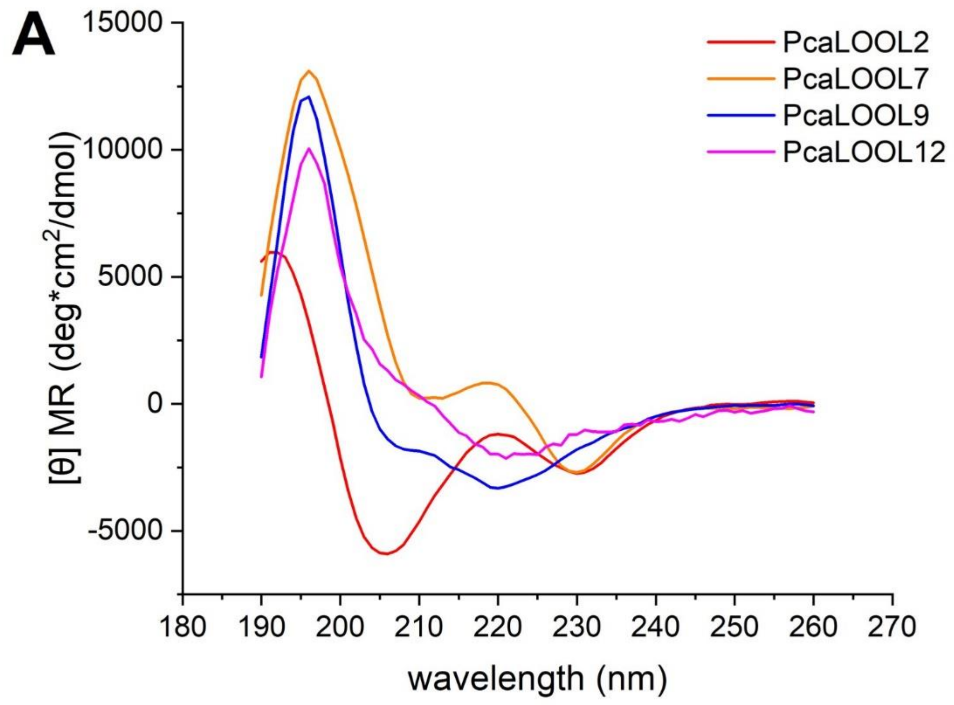


| B | PcaLOOL2 | PcaLOOL7 | PcaLOOL9 | PcaLOOL12 |
|---|---|---|---|---|
| Helix (%) | 9.9 | 3.3 | 8.5 | 2.1 |
| Anti parallel (%) | 28.9 | 35.2 | 28.4 | 36.7 |
| Parallel (%) | 0.0 | 1.9 | 0.0 | 0.0 |
| Turn (%) | 17.4 | 17.0 | 13.9 | 15.7 |
| Others (%) | 43.8 | 42.6 | 49.3 | 45.5 |
| Tm (°C) | 42.6 | 52.4 | 46.9 | 55.6 |

**Figure S3**

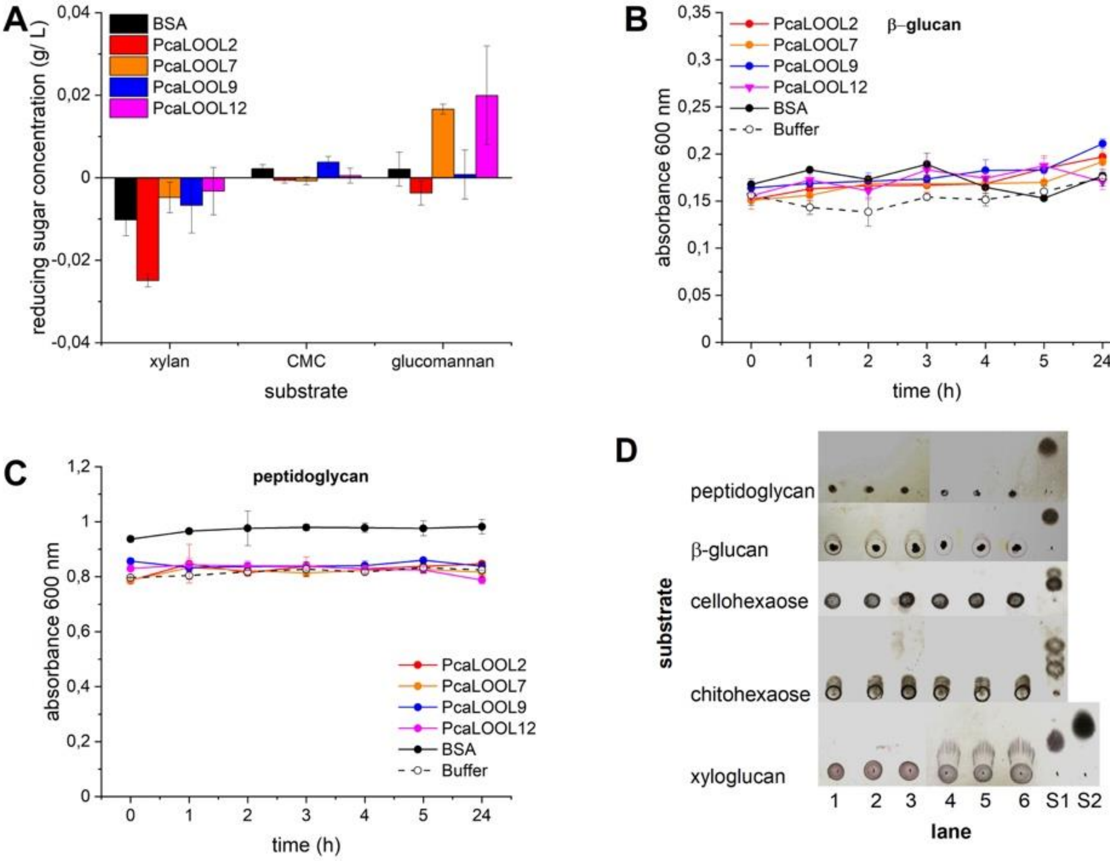
A
reducing sugar concentration (g/ L)
0,04
0,02
0
-0,02
-0,04
BSA
PcaLOOL2
PcaLOOL7
PcaLOOL9
PcaLOOL12
xylan
CMC
glucomannan
substrate
B
β-glucan
absorbance 600 nm
0,35
0,3
0,25
0,2
0,15
0,1
0,05
0
PcaLOOL2
PcaLOOL7
PcaLOOL9
PcaLOOL12
BSA
Buffer
0
1
2
3
4
5
24
time (h)
C
peptidoglycan
absorbance 600 nm
1,2
1
0,8
0,6
0,4
0,2
0
PcaLOOL2
PcaLOOL7
PcaLOOL9
PcaLOOL12
BSA
Buffer
0
1
2
3
4
5
24
time (h)
D
substrate
peptidoglycan
β-glucan
cellohexaose
chitohexaose
xyloglucan
1
2
3
4
5
6
S1
S2
lane

**Figure S4**

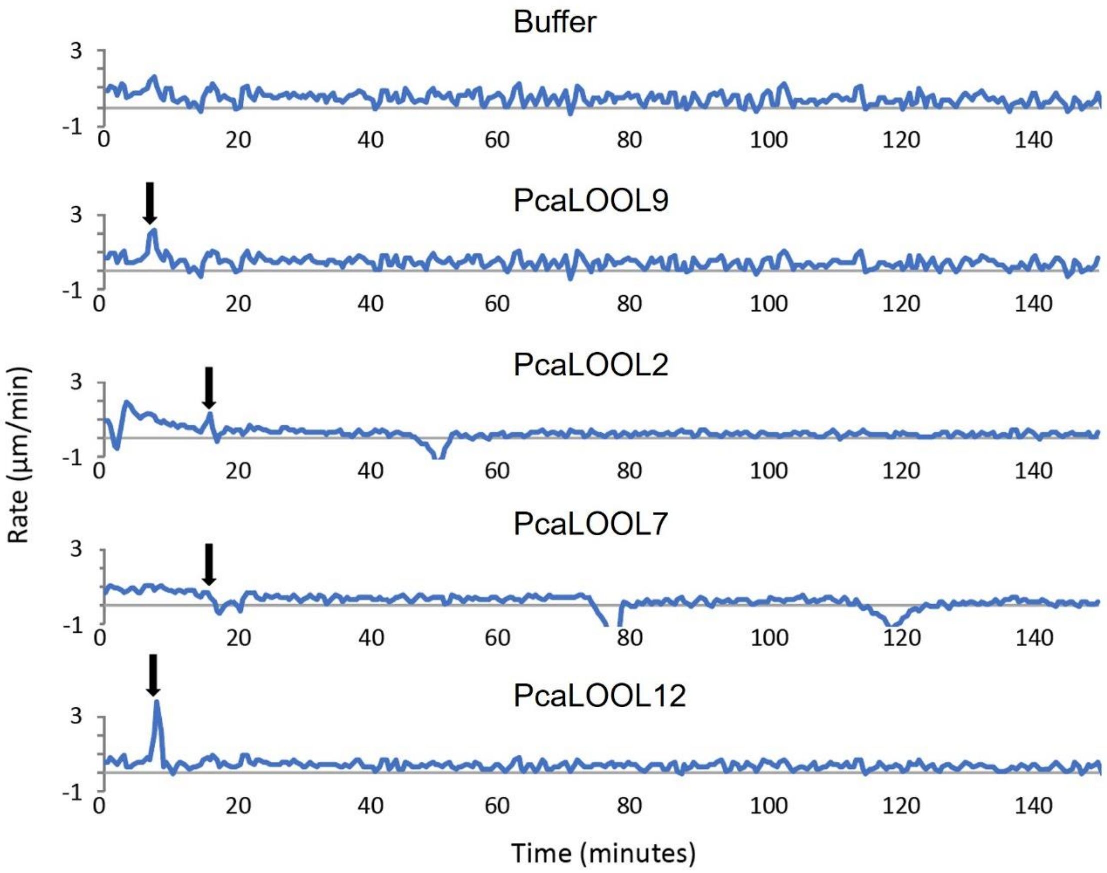
Buffer
PcaLOOL9
PcaLOOL2
PcaLOOL7
PcaLOOL12
Rate (µm/min)
Time (minutes)
3
-1
0
20
40
60
80
100
120
140

**Figure S5**

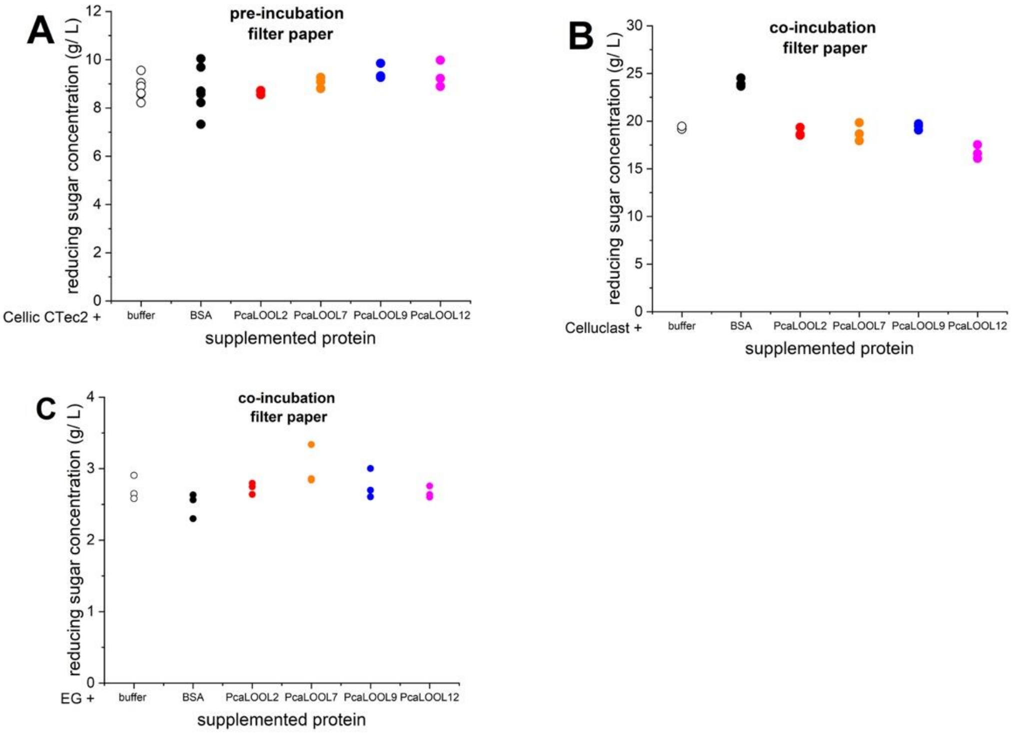
A
pre-incubation
filter paper
reducing sugar concentration (g/ L)
Cellic CTec2 +
buffer
BSA
PcaLOOL2
PcaLOOL7
PcaLOOL9
PcaLOOL12
supplemented protein
B
co-incubation
filter paper
reducing sugar concentration (g/ L)
Celluclast +
buffer
BSA
PcaLOOL2
PcaLOOL7
PcaLOOL9
PcaLOOL12
supplemented protein
C
co-incubation
filter paper
reducing sugar concentration (g/ L)
EG +
buffer
BSA
PcaLOOL2
PcaLOOL7
PcaLOOL9
PcaLOOL12
supplemented protein

**Figure S6**

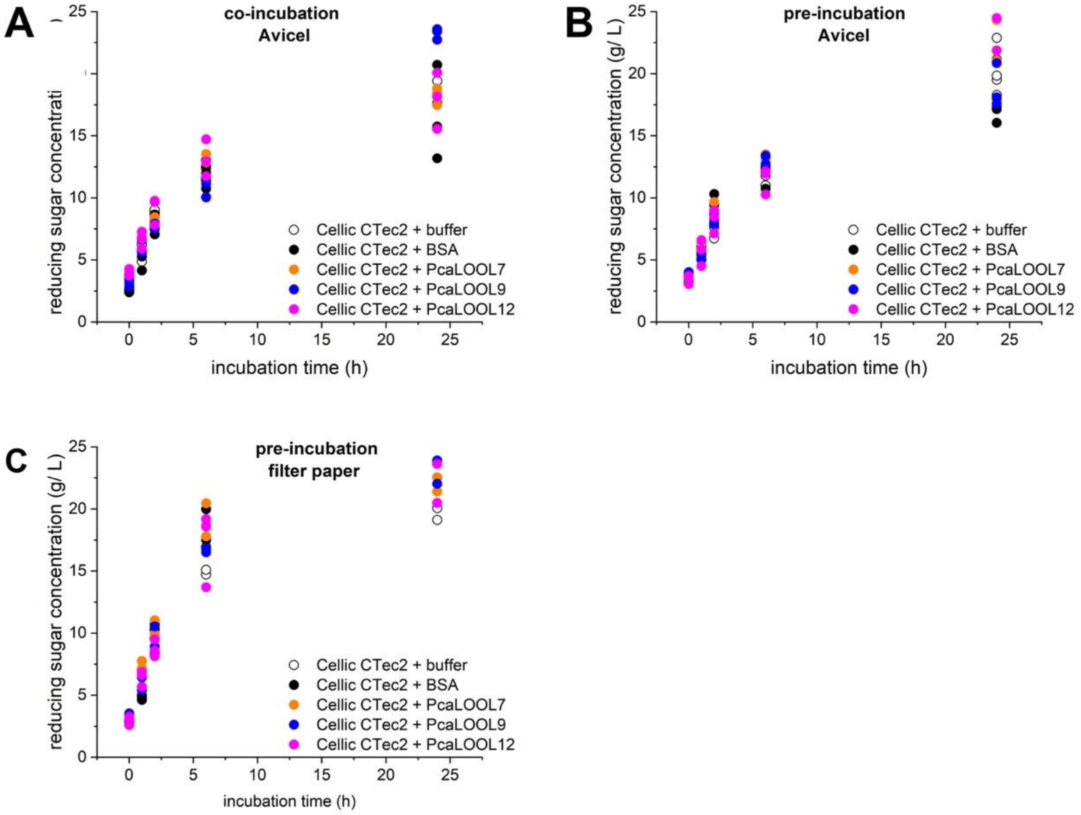


**Figure S7**

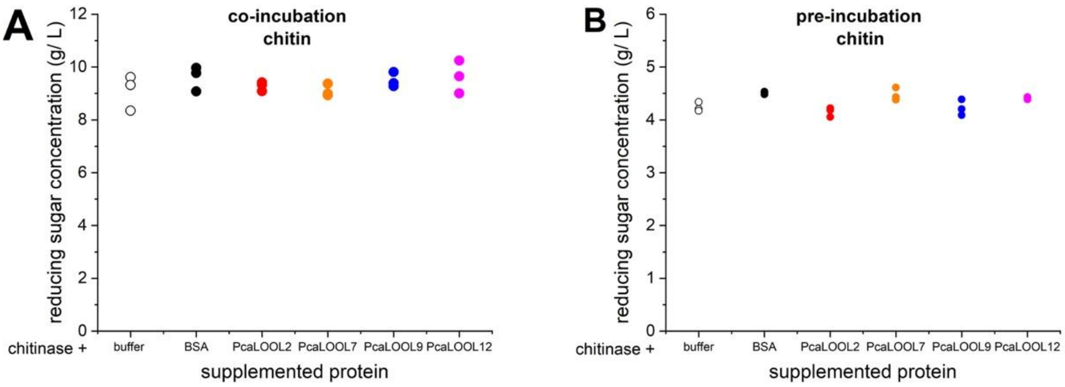


**Figure S8**

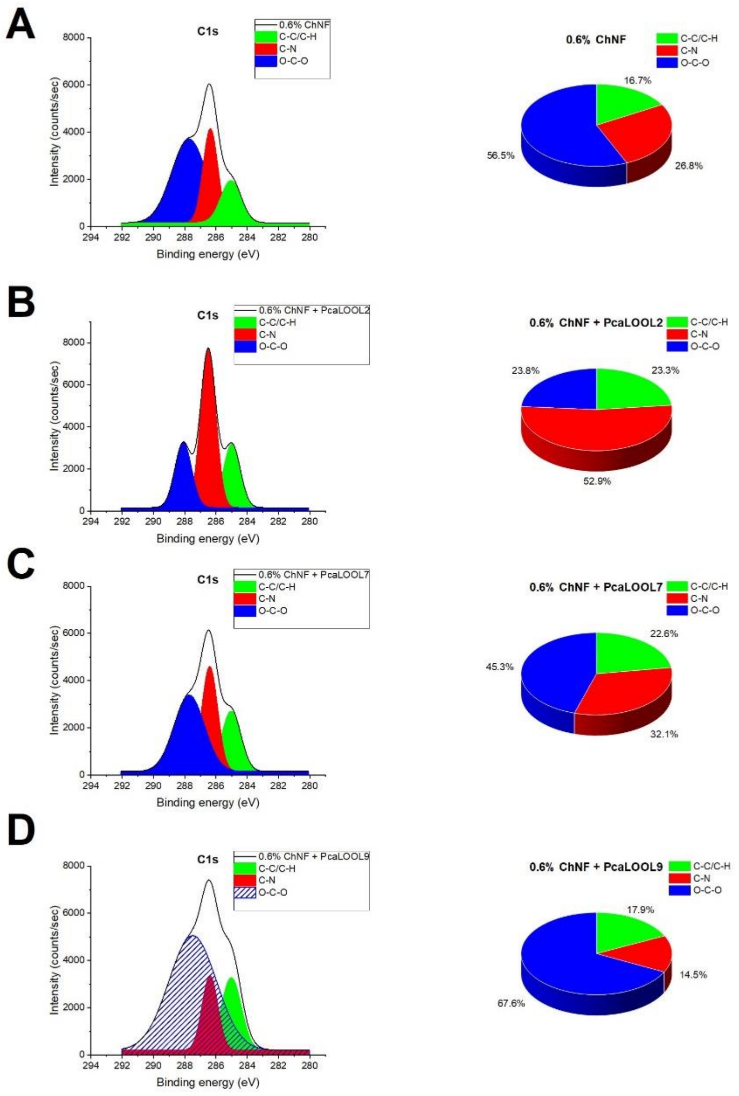
A
C1s
0.6% ChNF
C-C/C-H
C-N
O-C-O
Intensity (counts/sec)
Binding energy (eV)
0.6% ChNF
16.7%
26.8%
56.5%
B
C1s
0.6% ChNF + PcaLOOL2
C-C/C-H
C-N
O-C-O
Intensity (counts/sec)
Binding energy (eV)
0.6% ChNF + PcaLOOL2
23.8%
23.3%
52.9%
C
C1s
0.6% ChNF + PcaLOOL7
C-C/C-H
C-N
O-C-O
Intensity (counts/sec)
Binding energy (eV)
0.6% ChNF + PcaLOOL7
22.6%
45.3%
32.1%
D
C1s
0.6% ChNF + PcaLOOL9
C-C/C-H
C-N
O-C-O
Intensity (counts/sec)
Binding energy (eV)
0.6% ChNF + PcaLOOL9
17.9%
14.5%
67.6%

**Figure S9**
